# A feedback loop between the androgen receptor and 6-phosphogluoconate dehydrogenase (6PGD) drives prostate cancer growth

**DOI:** 10.1101/2020.09.02.279356

**Authors:** Joanna L. Gillis, Josephine A. Hinneh, Natalie Ryan, Swati Irani, Max Moldovan, Raj Shrestha, Lake Ee Quek, Andrew J. Hoy, Jeff Holst, Margaret M. Centenera, Ian G. Mills, David J. Lynn, Luke A. Selth, Lisa M. Butler

## Abstract

Alterations to androgen receptor (AR) signalling and cellular metabolism are hallmarks of prostate cancer. This study uncovers a novel link between AR and the pentose phosphate pathway (PPP) through 6-phosphogluoconate dehydrogenase (*6PGD*), an androgen-regulated gene that is upregulated in prostate cancer. Knockdown of 6PGD impairs growth and elicits death of prostate cancer cells, at least in part due to oxidative stress. Targeting 6PGD using 2 specific inhibitors, physcion and S3, was efficacious in multiple models of prostate cancer, including aggressive castration-resistant models. Importantly, S3 also suppressed proliferation of clinical patient-derived explants (PDEs). Mechanistically, 6PGD decreased expression and activity of AR in cell lines and PDEs, revealing a novel positive feedback loop between these factors. The enhanced efficacy of co-targeting AR and 6PGD further supported the biological relevance of this feedback. This work provides insight into the dysregulated metabolism of prostate cancer and supports investigation of co-targeting AR and the PPP.

## INTRODUCTION

Altered cellular metabolism is a hallmark of cancer. Perhaps the best characterised metabolic transformation in malignant cells is the so-called Warburg effect, in which cancer cells favour metabolism via glycolysis rather than the more efficient oxidative phosphorylation (1). While Warburg-like metabolism plays a key role in many malignancies, more recent work has demonstrated the diversity of cancer metabolism and revealed that tissue-of-origin is likely to be the critical determinant of malignant metabolic reprogramming (2). One tissue that exhibits a unique metabolic profile is the prostate (3). Normal prostate epithelial cells exhibit a truncated tricarboxylic acid (TCA) cycle to enable production of citrate, a key component of prostatic fluid, resulting in high rates of glycolysis (2). By contrast, malignant transformation switches metabolism of prostate cells to a more energetically favourable phenotype by re-establishing an intact TCA cycle, whereby citrate is utilised for oxidative phosphorylation and biosynthetic processes such as lipogenesis (4).

A major regulator of the unique metabolism of the normal and malignant prostate is the androgen receptor (AR) (5). AR is a hormone (androgen)-activated transcription factor that regulates expression of a large suite of genes involved in various aspects of metabolism, either directly or indirectly through activation of other master regulators such as SREBP (6, 7). Given its integral metabolic functions, it is unsurprising that AR is the primary oncogenic driver of prostate cancer (PCa) and the major therapeutic target in advanced and metastatic disease. While suppression of AR activity by androgen receptor pathway inhibitors (ARPIs) is initially effective in almost all men, prostate tumours inevitably develop resistance and progress to a lethal disease state known as castration-resistant prostate cancer (CRPC). One key feature of CRPC is the maintenance or re-activation of the AR signalling axis, as revealed by the therapeutic benefit of 2^nd^-generation ARPIs, such as the AR antagonist enzalutamide, in CRPC (8).

Unfortunately, the overall survival benefits of these newer ARPIs in men with CRPC are in the order of months (9), despite many tumours retaining dependence on AR (10). Collectively, these clinical observations highlight the ongoing dependence of CRPC on AR signalling and the intractable problems associated with therapies that inhibit this pathway.

Direct alterations to AR – including mutation, amplification, alternative splicing and altered ligand availability – have been well characterised as mechanisms of resistance in CRPC (11). However, the extent to which AR-mediated metabolic reprogramming is involved in therapy resistance in CRPC is less well understood. Herein, using an unbiased approach to discover potential PCa survival factors, we identify 6-phosphogluoconate dehydrogenase (6PGD) as a novel AR-regulated gene. 6PGD is a key enzyme in the phosphate pathway (PPP) (also referred to as the phosphogluconate pathway or the hexose monophosphate shunt), an alternative metabolic pathway for glucose breakdown. The PPP is comprised of two phases: an irreversible oxidative phase that generates NAPDH and ribulose-5-phosphate (Ru-5-P); and a subsequent reversible non-oxidative phase in which Ru-5-P is converted to R-5-P, a sugar precursor for generation of nucleotides (12). NADPH produced by the PPP is used for many anabolic reactions, including fatty acid synthesis, as well as an electron donor to generate reduced glutathione, the major endogenous antioxidant (13). Thus, the PPP is a major regulator of both redox homeostasis as well as anabolic reactions, depending on cellular requirements. We demonstrate that 6PGD plays a key role in PCa growth and survival, at least in part through moderating oxidative stress, and uncover a novel feedback mechanism linking 6PGD and the AR signalling axis that provides impetus for further investigation of co-targeting AR and the PPP as a novel therapeutic strategy.

## RESULTS

### *6PGD* is an androgen-regulated gene in prostate cancer

The current clinical ARPIs, such as enzalutamide, do not target the entire repertoire of genes regulated by the AR in prostate tumour cells (14). We hypothesised that ablation of AR expression would be the most appropriate “therapeutic benchmark” to identify the key regulators of tumour cell survival regulated by AR. To qualitatively and quantitatively compare downstream responses to AR ablation and AR antagonism, LNCaP cells were treated with AR siRNA (siAR; i.e. AR ablation) or enzalutamide (Enz; AR antagonism) and subsequently evaluated by RNA-seq. The experimental conditions were optimised to achieve comparable suppression of the canonical AR target, PSA, which is encoded by the *KLK3* gene (Figure 1A). Genes affected by siAR were highly concordant with an independent dataset (15) (Figure S1). As expected, most (78 %) genes altered by enzalutamide (compared to vehicle control) were also similarly dysregulated by siAR (compared to a control siRNA, siCon) (Figure 1B; Dataset S1). An additional 2,574 genes were altered in their expression by siAR but not enzalutamide (Figure 1B, q < 0.05). On closer examination, many of these genes were altered in their expression by enzalutamide but not sufficiently for them to be identified as statistically significant differentially expressed genes. A further direct statistical comparison of gene expression between the two treatment groups identified that there were 581 genes that were differentially expressed in the siAR treated cells compared to those treated with enzalutamide including, as expected, *AR* itself (Figure 1B-C, Dataset S1). These results provide further evidence for the hypothesis that AR ablation is more effective at suppressing the AR-regulated transcriptome compared with AR antagonism, at least in this experimental system.

**Figure 1.**
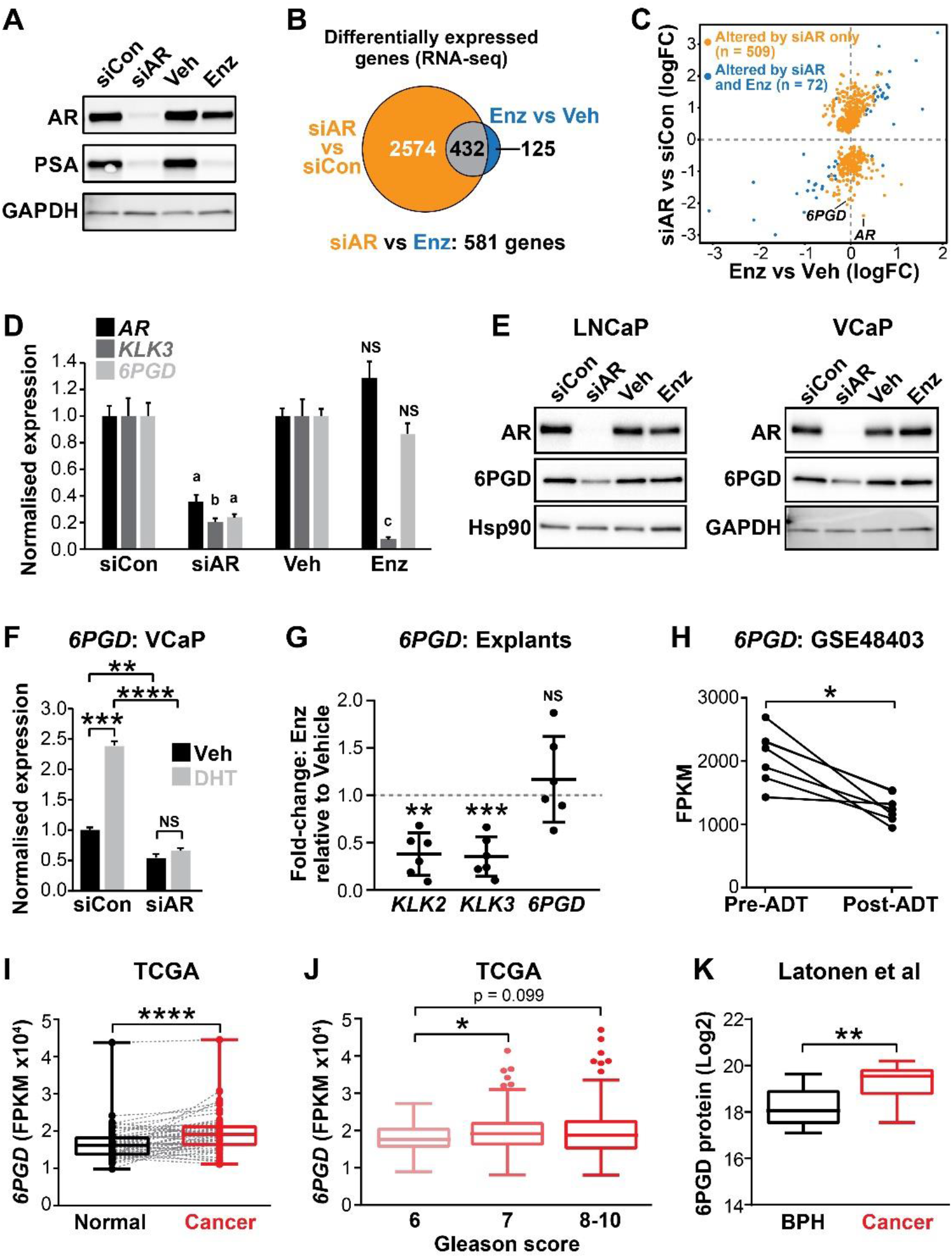
6PGD is an AR-regulated gene and is elevated in prostate cancer. **(A)** Effect of siAR and enzalutamide (Enz) on the AR target, PSA. LNCaP cells were transfected with AR (siAR; 12.5 nM) or control (siCon) siRNA for 48 h or treated with Enz (1 µM) or vehicle (Veh) for 24 h, after which AR and PSA proteins were evaluated by immunoblotting. GAPDH was used as loading control. **(B)** Numbers of genes differentially expressed by siAR (vs siCon) or Enz (vs Veh) are shown in the Venn diagram (at top). Below: an alternative analysis identified 581 genes differentially expressed by siAR versus Enz. **(C)** Scatterplot of genes affected by siAR and Enz. The 581 genes differentially expressed by siAR versus Enz are shown in blue (n = 72, genes differentially expressed by siAR versus siCon and Enz versus Veh) and yellow (n = 509, genes differentially expressed by siAR versus siCon but not by Enz versus Veh. **(D)** Validation of *6PGD* expression in response to siAR and Enz by RT-qPCR. Gene expression was normalised to *GUSB* and *L19* and represents the mean ± standard error (SE) of three biological replicates; siCon and Veh were set to 1. Differential expression was evaluated using unpaired t tests (a, p < 0.01; b, p < 0.001; c, p < 0.0001; NS, not significant). **(E)** 6PGD protein levels in response to siAR and Enz treatments were measured by immunoblotting in LNCaP (left) and VCaP (right) cells. HSP90 and GAPDH were used as loading controls. **(F)** RT-qPCR of *6PGD* expression in response to DHT and siAR in VCaP cells. Cells were transfected with siRNAs for 24 h, and then treated with 1 nM DHT for another 24 h. Gene expression was normalised and graphed as in D. Differential expression was evaluated by t tests (*, p < 0.05). **(G)** RT-qPCR of *KLK2, KLK3* and *6PGD* expression in response to Enz treatment (1 µM, 72 h) in patient-derived explants. Gene expression was normalised to *GAPDH, PPIA* and *TUBA1B* and is represented as fold-change of enzalutamide relative to vehicle treatment. Differential expression was evaluated by one sample t tests (**, p < 0.01; ***, p < 0.001). **(H)** *6PGD* mRNA expression in prostate tumours (GSE48403). A Wilcoxon matched-pairs signed rank test was used to compare expression in the groups. FPKM, fragments per kilobase of exon per million mapped reads. **(I)** *6PGD* expression is elevated in primary prostate cancer. The TCGA dataset comprises 52 patient-matched normal and cancer samples. Boxes show minimum and maximum (bottom and top lines, respectively) and mean (line within the boxes) values. A paired t test was used to compare expression in normal versus cancer. FPKM, fragments per kilobase of exon per million mapped reads. **(J)** *6PGD* expression by Gleason grade in the TCGA cohort. Boxes show minimum and maximum (bottom and top lines, respectively) and mean (line within the boxes) values. Unpaired t tests were used to compare expression between the groups. FPKM, fragments per kilobase of exon per million mapped reads. **(K)** 6PGD protein expression in clinical prostate samples (benign prostatic hyperplasia (BPH) and tumours) were measured my mass spectrometry. Boxes show minimum and maximum (bottom and top lines, respectively) and mean (line within the boxes) values. An unpaired t test was used to compare expression between the groups.

The gene most significantly associated with AR ablation and not AR antagonism was *6PGD* (Figure 1C, Dataset S1), which encodes an enzyme in the pentose phosphate pathway (PPP). We confirmed that 6PGD expression was down-regulated by AR knockdown but not by acute AR antagonism in multiple PCa cell lines (LNCaP and VCaP) at both the mRNA and protein level (Figure 1D-E, Figure S2). Down-regulation of 6PGD was also seen with a second AR siRNA, validating 6PGD as a *bona fide* target of AR (Figure S2). In further support of differential regulation by siAR versus AR antagonism, neither of the newest clinically approved AR antagonists (apalutamide and darolutamide) altered 6PGD protein or mRNA expression (Figure S3). Conversely, AR activation with the androgen 5α-dihydrotestosterone (DHT) stimulated *6PGD* expression, and this effect was abolished by co-treatment with siAR (Figure 1F). To determine whether AR inhibition affects 6PGD in more biologically relevant systems, we first utilised our patient-derived explant (PDE) model (16). Similar to 2-dimensional PCa cell line culture, we did not observe enzalutamide-mediated changes to *6PGD* mRNA expression in the PDE model over a time-frame of 48h, under conditions that caused significant repression of the well-characterised AR target genes *KLK2* and *KLK3* (Figure 1G). By contrast, longer term (∼14 weeks) androgen deprivation therapy in patients caused a significant decrease in *6PGD* mRNA levels (Figure 1H). Collectively, these findings reveal 6PGD as a novel AR-regulated factor in both PCa cell lines and clinical samples.

As an initial assessment of the relevance of 6PGD in clinical PCa, we examined its expression in the TCGA dataset (17) and found that *6PGD* mRNA expression was significantly elevated in cancer compared to patient-matched normal tissue and also showed an association with increasing Gleason grade (Figure 1I-J). An association with malignancy was recapitulated at the protein level (Figure 1K) in a distinct set of patient samples for which proteomes were profiled using mass spectrometry (18). We further examined 6PGD protein expression in prostate tumours by immunohistochemistry (IHC). 6PGD was detected in all tissues that were examined and was predominantly localised to the cytoplasm and peri-nuclear regions of epithelial cells (Figure S4). Moreover, we observed a trend towards increasing protein levels in the more aggressive tumours (Figure S4). In summary, 6PGD is highly expressed in prostate tumours, suggesting that the PPP may play an important metabolic role in this cancer type.

### SREBP mediates induction of *6PGD* downstream of the androgen receptor

AR binds to gene enhancers or promoters to directly regulate transcription (19). However, we found no clear evidence of AR binding sites proximal the *6PGD* transcriptional start site in genome-wide DNA binding (ChIP-seq) datasets from tissues and cell lines (Figure 2A and data not shown), suggesting that the AR pathway may indirectly regulate *6PGD* expression via another downstream pathway(s) or factor(s). One credible intermediary between AR and 6PGD is sterol regulatory element-binding protein-1 (SREBP1), a transcriptional master regulator of genes with a role in lipid and cholesterol production (20). AR enhances SREBP1 expression and activity in a multifaceted manner, most notably by upregulating the SREBP1 activator SCAP (20) and by activating the mTOR pathway, which in turn leads to elevated SREBP1 expression (21). Additionally, SREBP1 has been proposed to directly regulate *6PGD* in mouse adipocytes by direct binding to its promoter (22). We mined ENCODE SREBP1 ChIP-seq data and identified an SREBP1 binding site at the *6PGD* promoter in two cancer cell lines, HEPG2 (liver) and MCF7 (breast) (Figure 2B). Regulation of 6PGD by SREBP1 in prostate cancer cells was confirmed by siRNA-mediated knockdown of SREBP1 (Figure 2C). To test whether SREBP1 acts downstream of AR to increase *6PGD* expression, we treated cells with a combination of DHT and Fatostatin, an inhibitor of SREBP1, and found Fatostatin to effectively suppress DHT-mediated induction of *6PGD* (Figure 2D). Collectively, these results are indicative of an AR-SREBP1-6PGD circuit in prostate cancer cells and implicate SREBP1 as a key mediator of PPP activation by AR.

**Figure 2.**
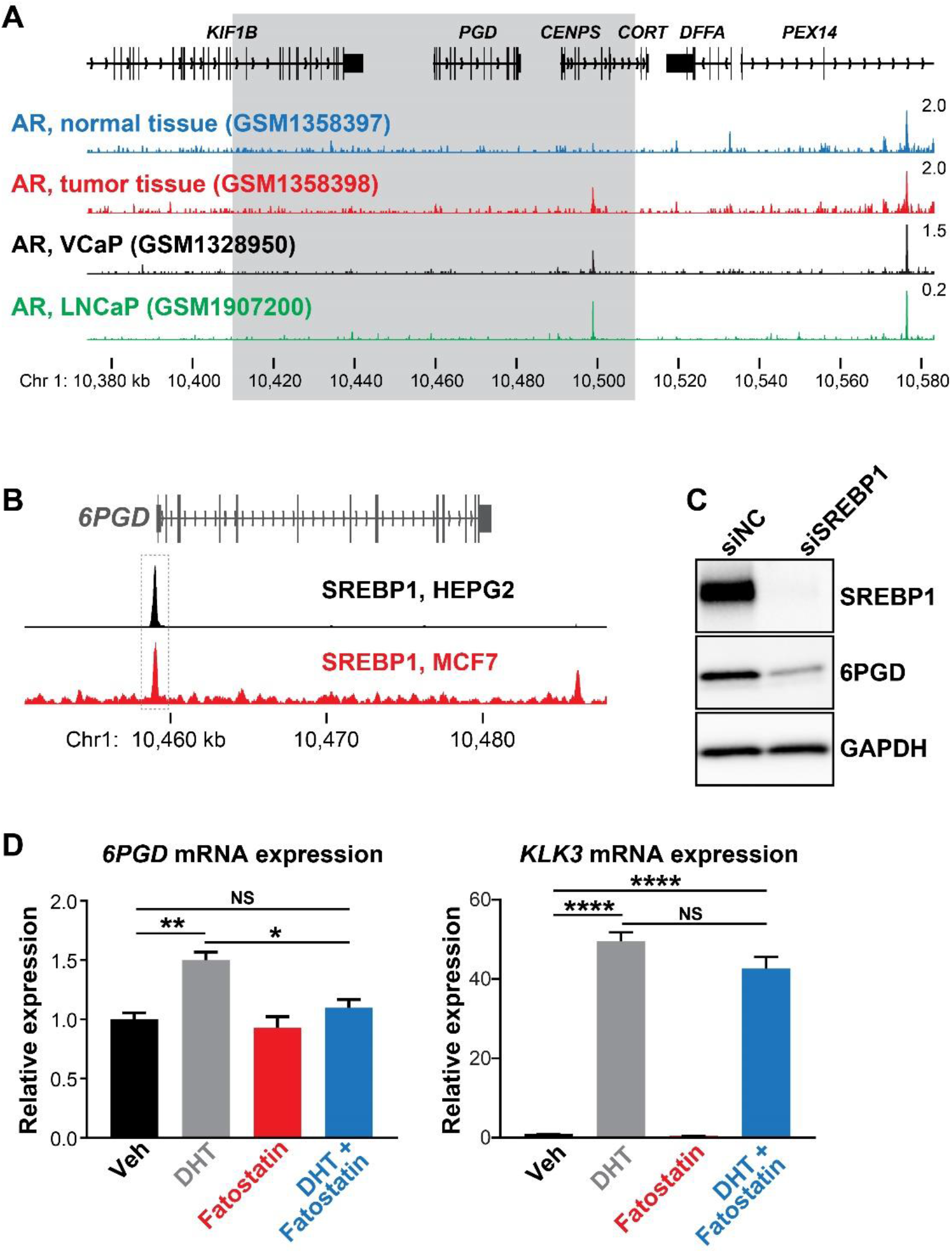
**(A)** ChIP-seq data showing AR DNA binding near the *6PGD* gene in non-malignant and prostate tumor samples (23) and the LNCaP (24) and VCaP (14) cell line models. The grey box indicates a region +/-50kb of the *6PGD* transcriptional start site. **(B)** ChIP-seq data showing SREBP1 DNA binding at the *6PGD* promoter in HEPG2 and MCF7 cells. Data is from ENCODE (25) (HEPG2: ENCFF000XXR; MCF7: ENCFF911YFI). **(C)** Effect of siSREBP1 on 6PGD protein. LNCaP cells were transfected with siRNA (siSREBP1; 12.5 nM) or control (siCon) for 72 h after which SREBP1 and 6PGD protein levels were evaluated by immunoblotting. GAPDH was used as loading control. **(D)** RT-qPCR of *6PGD* expression in response to DHT and Fatostatin in LNCaP cells. Cells were serum starved in charcoal-stripped FBS media for 72 h, and then treated with Veh or 10 nM DHT +/-10 µM Fatostatin for another 24 h. Gene expression was normalised to *GUSB* and *L19* and represents the mean ± standard error (SE) of three biological replicates. Differential expression was evaluated by t tests (*, p < 0.05; **, p < 0.01).

### 6PGD is required for the growth of prostate cancer cells and has downstream effects on AMPK and ACC1 activity

Regulation of 6PGD by the AR signalling axis supports other recent reports linking the PPP to PCa (13, 26); and although the role of the PPP in this malignancy is not fully elucidated, it could serve to fuel cell growth and protect against oxidative stress. In support of this, knockdown of 6PGD with two highly effective siRNAs (Figure S5) significantly decreased viability (Figure 3A) and increased death (Figure 3B) of LNCaP and VCaP cells. Concomitant with these phenotypic effects, mass spectrometry revealed accumulation of 6PGD’s substrate, 6-phosphogluconate (6-PG) (Figure 3C), confirming specificity of the knockdown. Since a key role of the PPP is to regulate intracellular redox state, we also measured ROS using a flow cytometric-based assay. As expected, knockdown of 6PGD (and AR) significantly increased levels of intracellular ROS (Figure 3D), which could be reversed by the antioxidant Trolox (Figure 3E).

**Figure 3.**
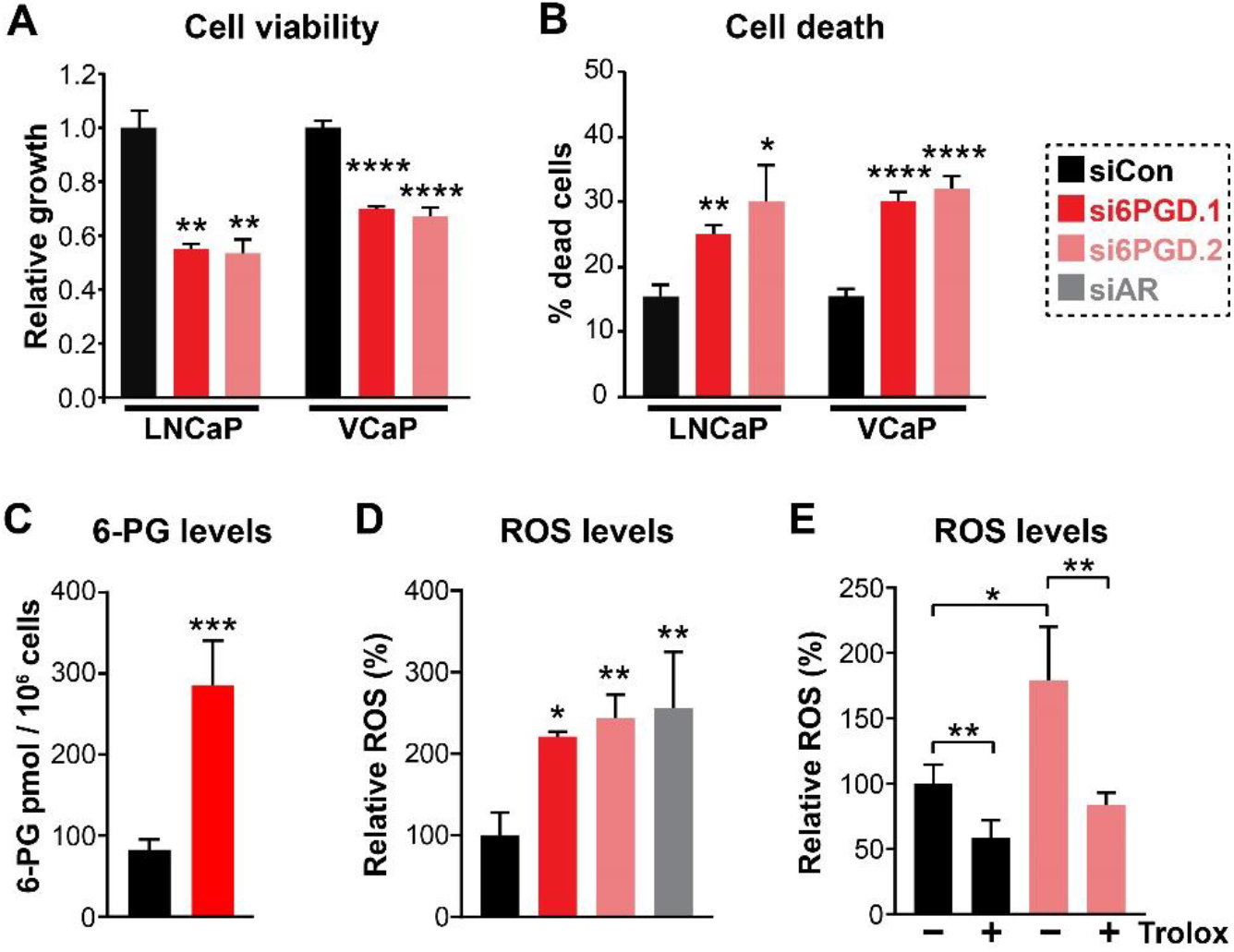
Knockdown of 6PGD has multi-faceted anti-cancer effects in prostate cancer cells. **(A-B)** Knockdown of 6PGD with two distinct siRNAs (si6PGD.1 and si6PGD.2) reduced LNCaP viability (A) and increased cell death (B), as assessed using Trypan blue exclusion assays. Bars are mean ± SE of triplicate samples, and are representative of 3 independent experiments. Effects were evaluated using t tests (*, p < 0.05; **, p < 0.01). **(C)** Knockdown of 6PGD causes accumulation of intracellular 6-PG in LNCaP cells, as determined by mass spectrometry. Results are representative of 2 independent experiments. Effects were evaluated using t tests (p < 0.001). **(D)** Knockdown of 6PGD and AR causes increased levels of reactive oxygen species (ROS) in LNCaP cells. Data was normalised to siCon, which was set to 100%. Effects were evaluated using t tests (*, p < 0.05; **, p < 0.01). **(E)** ROS production in response to si6PGD is rescued by the antioxidant Trolox. Data is presented as in C.

### Inhibition of 6PGD suppresses prostate cancer growth and increases ROS

Having established that 6PGD is required for optimal PCa cell growth and protects against oxidative stress, we evaluated pharmacological targeting of this enzyme as a potential therapeutic strategy. Physcion, a plant-derived anthraquinone, was recently identified as an inhibitor of 6PGD using an *in vitro* screening assay (27). Treatment of LNCaP cells with physcion dose-dependently inhibited growth and elicited death (Figure S6). However, low solubility limits the pre-clinical and clinical utility of this compound. Therefore, we focussed our efforts on a derivative of physcion, S3, which has substantially improved solubility (∼50-fold: 1 mM physcion c.f. 50 mM S3 in DMSO) (27). Similarly to physcion, S3 reduced LNCaP cell viability and caused cell death (Figure 4A-B). Cell kill was at least partly mediated via apoptosis, as demonstrated by flow cytometric-based Annexin/7-AAD assay (Figure 4C). Importantly, S3 increased levels of cellular ROS in a dose-dependent manner (Figure 4D), strengthening the link between the PPP and control of redox homeostasis. S3 was active in a range of PCa models, including VCaP and models of CRPC (V16D and MR49F) (Figure 4E-F). The efficacy of S3 in MR49F cells was particularly notable, since this aggressive LNCaP-derived line is resistant to the 2^nd^-generation AR antagonist enzalutamide (28). S3 was also growth inhibitory in AR-negative PC3 cells, although this line was less sensitive than AR-driven models (Figure S7). To assess the potential of targeting 6PGD with S3 in a more clinically-relevant setting, we exploited the PDE model (16). Notably, S3 reduced proliferation, as measured by IHC for Ki67, in all tumours (n = 9) that were evaluated (Figure 4G).

**Figure 4.**
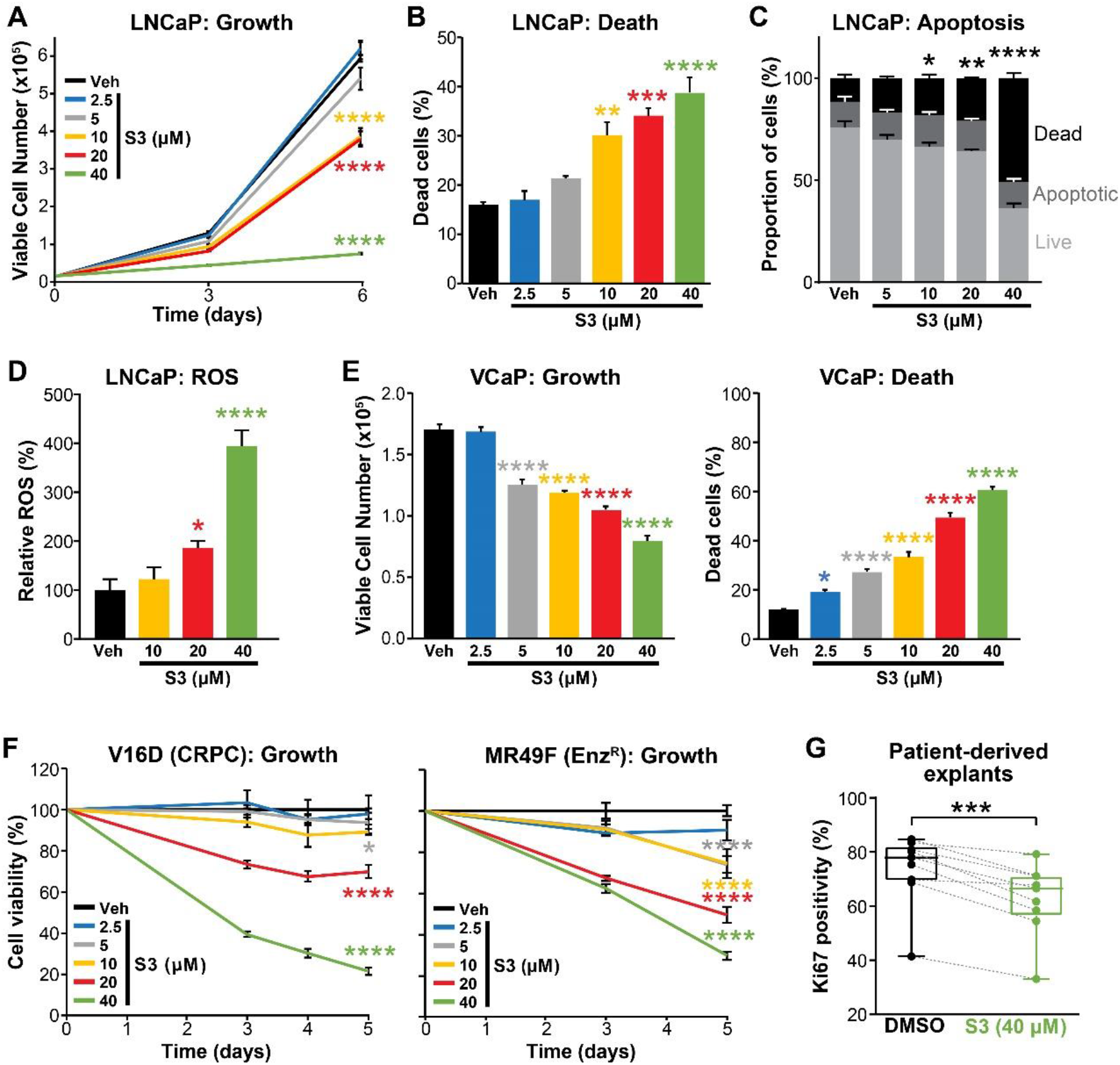
Pharmacological targeting of 6PGD in prostate cancer. **(A-B)** The 6PGD inhibitor, S3, dose dependently decreased viability (A) and increased death (B) of LNCaP cells, as determined by Trypan blue exclusion assays. Dead cells were counted at day 6. Data represents the mean ± SE of triplicate samples and are representative of 3 independent experiments. Growth (day 6) and death for each dose was compared to vehicle using ANOVA and Dunnett’s multiple comparison tests (****, p < 0.0001). Veh, vehicle. **(C)** S3 causes apoptosis of LNCaP cells, as determined using flow cytometry-based Annexin V/7-AAD assays. Cells were assessed 72 h after treatment. Data represents the mean ± SE of triplicate samples and are representative of 4 independent experiments. Dead cell proportions were compared to vehicle using ANOVA and Dunnett’s multiple comparison tests (*, p < 0.05; **, p < 0.01; ****, p < 0.0001). **(D)** S3 causes increased levels of reactive oxygen species (ROS) in LNCaP cells. Data was normalised to Veh, which was set to 100%. Effects were evaluated using t tests (*, p < 0.05; ****, p < 0.0001). **(E)** S3 dose dependently decreased viability (left) and increased death (right) of VCaP cells, as determined by Trypan blue exclusion assays. Live and dead cells were counted 4 days after treatment. Data represents the mean ± SE of triplicate samples and are representative of 3 independent experiments. **(F)** S3 suppresses the growth of CRPC cells (V16D) and enzalutamide-resistant CRPC cells (MR49F), as determined using CyQuant Direct Cell Proliferation Assay. Fluorescence from day 0 was set to 100%. Data represents the mean ± SE of triplicate samples and are representative of 2 independent experiments. **(G)** S3 inhibits the proliferation of prospectively collected human tumours grown as patient-derived explants (PDEs). PDEs (from n = 9 patients) were treated for 72 h. Ki67 positivity, a marker of proliferation, was determined using IHC. Boxes show minimum and maximum (bottom and top lines, respectively) and mean (line within the boxes) values. A paired t test was used to compare Ki67 positivity in treated versus control samples (***, p < 0.001).

In addition to directly promoting cell growth and survival via anabolism and limiting oxidative stress, the PPP has been reported to suppress AMPK activity by inhibiting its phosphorylation (29), thereby activating key anabolic pathways via acetyl-CoA carboxylase 1 (ACC1) and mammalian target of rapamycin (mTORC1) (Figure 5A). Accordingly, we examined whether these pathways are altered in PCa cells by pharmacological targeting of 6PGD. S3 treatment activated AMPK and repressed ACC1 and mTOR pathways in a dose-dependent manner in multiple PCa cell lines, as revealed by increased levels of phospho-AMPK (pAMPK) and phospho-ACC1 (pACC1) and decreased levels of phospho-S6K (pS6K) / phospho-S6 (pS6) (Figure 5B-C). Importantly, we recapitulated the impact of S3 on mTOR signalling in our tumour PDE system (Figure 5C). Collectively, these results reveal that PPP is an upstream regulator of AMPK, ACC1 and mTOR in prostate cancer, a key implication being that targeting 6PGD could impede multiple cancer-promoting metabolic pathways.

**Figure 5.**
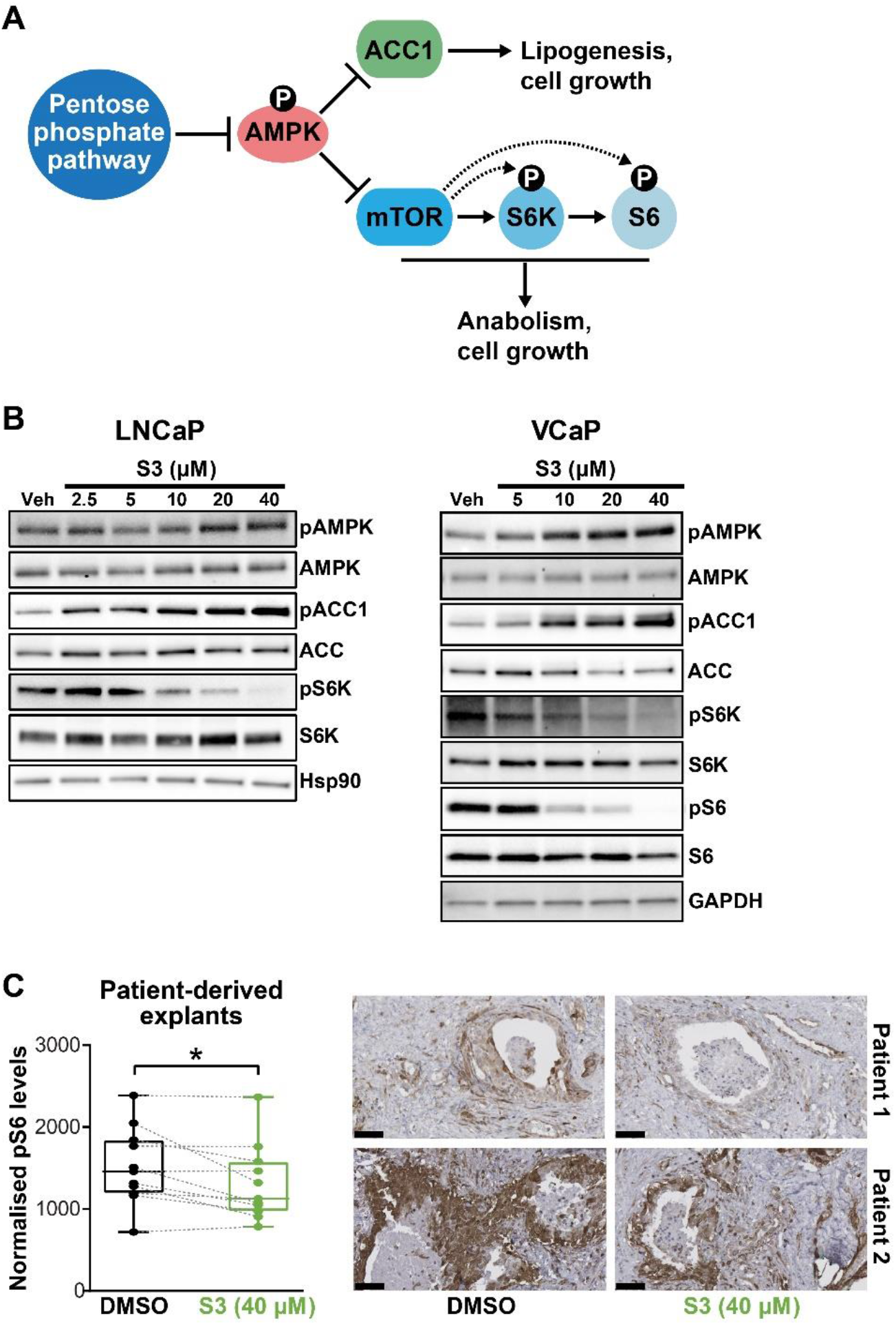
Targeting 6PGD activates AMPK and represses ACC1 and mTOR pathways. **(A)** Schematic showing key metabolic pathways downstream of the PPP. By suppressing AMPK signalling, the PPP can enhance the activity of ACC1 and mTOR and subsequently various growth-promoting anabolic processes. **(B)** S3 activates AMPK and inhibits ACC1 and mTOR signalling, as determined by increased levels of pAMPK and pACC1, respectively. LNCaP (left) and VCaP (right) cells were treated for 24 h with the indicated doses of S3 prior to analysis of proteins by immunoblotting. **(C)** S3 inhibits mTOR signalling, as indicated by reduced pS6, in patient-derived explants (PDEs). PDEs (from n = 11 patients) were treated for 72 h. The levels of pS6 were measured using IHC. Boxes (graph on left) show minimum and maximum (bottom and top lines, respectively) and mean (line within the boxes) values. A paired t test was used to compare Ki67 positivity in treated versus control samples (***, p < 0.001). Representative IHC images are shown on the right (scale bars represent 50 µm).

### A feedback loop between AR and 6PGD supports combinatorial targeting of these factors

During our investigations into the mode of action of S3 and physcion, we noted that both agents reduced steady-state levels of AR protein in models of castration-sensitive and castration-resistant prostate cancer (Figure 6A; Figure S8A). This observation suggested that targeting 6PGD would inhibit the AR signalling axis. We validated this hypothesis by demonstrating that S3 and physcion dose dependently reduced the expression of AR target genes in multiple cell line models (Figures 6A-B, Figure S8B-D) and, critically, in our clinical PDE tissues (Figure 6C). These observations reveal a positive feedback loop involving 6PGD and the AR signalling axis, and hence suggest co-targeting of AR and 6PGD as a rational combination therapy. In support of such an approach, enzalutamide and S3 exhibited an additive effect in VCaP cells for growth inhibition (Figure 6D) and induction of cell death (Figure 6E) compared to the single agents. The value of such a combinatorial targeting strategy was further validated using a cell line model of CRPC (Figure 6F).

**Figure 6.**
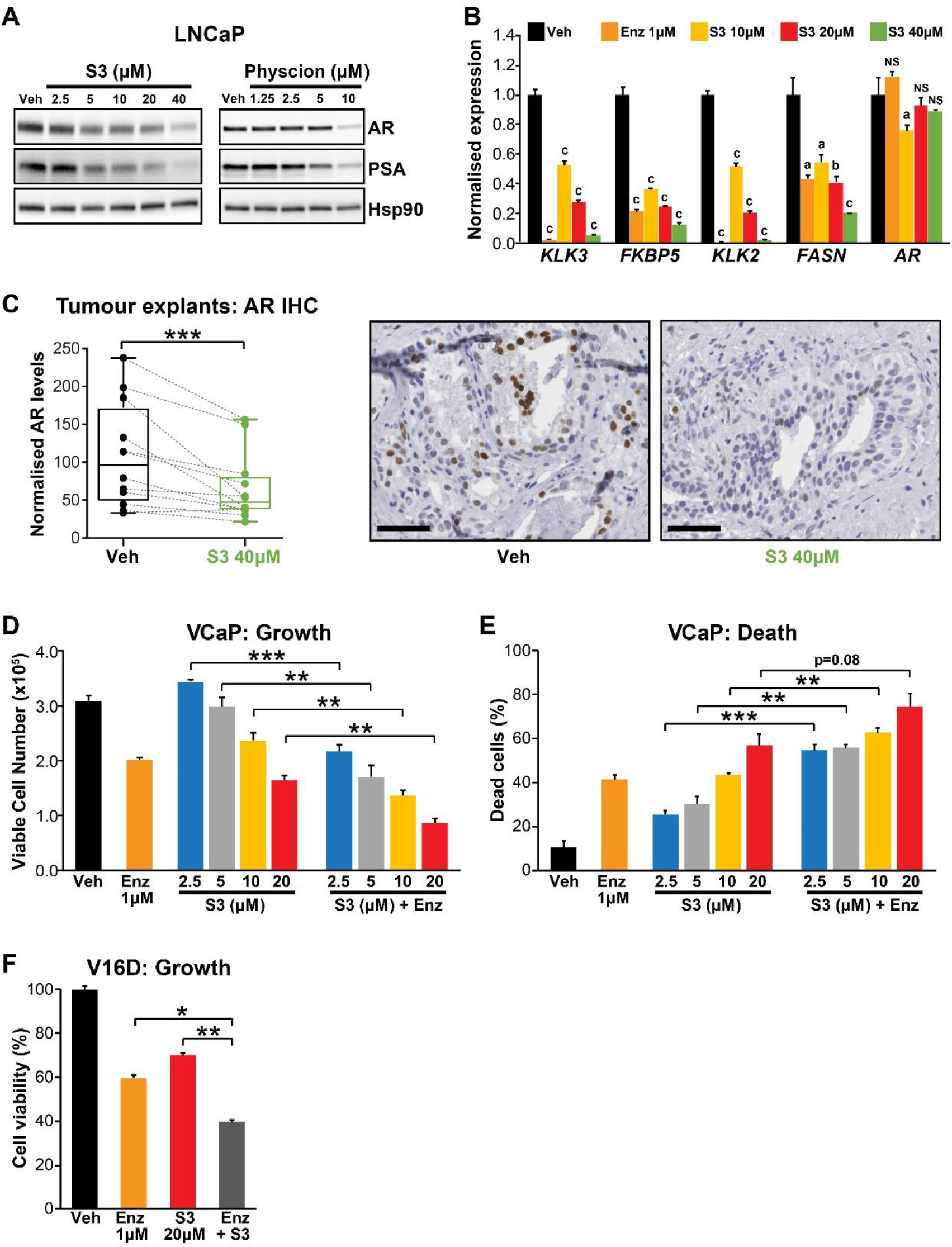
Targeting the AR/PGD feedback loop in prostate cancer. **(A)** Protein levels of AR and its target in response to S3 (24 h of treatment) and physcion (48 h of treatment) in LNCaP cells, as determined by immunoblotting. HSP90 was used as a loading control. **(B)** AR target gene expression in response to S3 treatment in LNCaP cells, as determined by RT-qPCR. Gene expression was normalised to *GUSB* and *L19* and represents the mean ± standard error (SE) of three biological replicates; Veh was set to 1. Differential expression was evaluated using ANOVA and Dunnett’s multiple comparison tests (a, p < 0.01; b, p < 0.001; c, p < 0.0001; NS, not significant). **(C)** S3 reduces AR protein levels in PDEs. AR levels in tumours from 14 patients were measured by IHC (left). Boxes show minimum and maximum (bottom and top lines, respectively) and mean (line within the boxes) values. A paired t test was used to compare AR levels in treated versus control samples (***, p < 0.001). Representative IHC images are shown on the right (scale bars represent 50 µm). **(D-E)** Anti-cancer effects of combined Enz and S3 treatment in VCaP cells. Live (D) and dead (E) cells were measured by Trypan blue exclusion assays 4 days after treatment. Data represents the mean ± SE of triplicate samples and are representative of 3 independent experiments. **(F)** Anti-cancer effects of combined Enz and S3 treatment in V16D cells. Live cells (F) were measured as in D after 3 days of treatment; data are representative of 3 independent experiments.

## DISCUSSION

Prostate cancer possesses a unique androgen-regulated metabolic profile, characterised by high rates of lipogenesis and oxidative phosphorylation compared to the normal state. More recently, altered glucose metabolism has emerged as another feature of this common malignancy (3). In this study, we identified *6PGD* as an AR-regulated gene that may not be effectively suppressed in tumour cells by current ARPIs such as enzalutamide. 6PGD is the third enzyme in a critically important glucose metabolic pathway, the PPP. Our data reveal that a positive feedback loop between AR and 6PGD promotes growth and survival of tumour cells. This work not only expands our knowledge of the interplay between hormones and glucose metabolism in PCa but also exposes a new therapeutic vulnerability.

Our identification of 6PGD as an androgen-regulated PPP enzyme lends further support to this pathway being a key metabolic target of androgens in prostate cancer. Frigo and colleagues recently demonstrated that G6PD, the rate-limiting enzyme of this pathway, is also transcriptionally and post-transcriptionally regulated by AR signalling (13). Moreover, an enzyme that regulates the non-oxidative phase of the PPP, transketolase-like protein 1 (TKTL1), increases in expression during PCa progression, being highest in metastatic tumours (30). Such multi-level control of a single pathway emphasises the relevance of increased PPP flux in PCa. It is notable that the androgen-regulated enzymes of this pathway, 6PGD and G6PD, both catalyse steps in the NADPH-generating oxidative phase of the PPP; this represents another mechanism underlying hormonal protection against oxidative stress in the prostate.

Despite its role as a key downstream effector of androgen-regulated cellular metabolism, our data do not support a direct mode of transcriptional regulation of *6PGD* by AR. Rather, AR harnesses another key metabolic transcription factor, SREBP1, to drive expression of 6PGD and hence activity of the PPP. SREBP1 is a transcription factor that regulates genes involved in fatty acid and cholesterol biosynthesis and homeostasis, and itself a therapeutic target in prostate cancer (31). Although some metabolic genes appear to be directly co-regulated by AR and SREBP1 based on the binding of both factors to cis-regulatory elements (e.g. *FASN*, (32, 33)), our observation that fatostatin largely blocks androgen-mediated induction of *6PGD* supports an indirect role for AR’s transcriptional regulatory function in this process. A broader implication of the AR-SREBP1-6PGD circuit identified in this study is its potential relevance as a clinical target; therapeutic strategies that effectively suppress this circuit would impinge on the activity of 3 important and distinct oncogenic drivers.

We propose that AR-mediated activation of the PPP in PCa would yield additional advantages beyond the generation of key substrates for nucleic acid anabolism and the antioxidant NADPH. Most notably, PPP suppression of AMPK, itself a hub for cellular metabolic and growth control, results in augmentation of ACC1 and mTOR activity (34). The importance of both ACC1 and mTOR in enabling PCa cells to meet their energy demands is increasingly well recognised; indeed, both of these factors are key mediators of *de novo* lipogenesis, high levels of which are a fundamental attribute of prostate tumours (35). Mechanistically, it has been reported that 6PGD-mediated production of Ru-5-P inhibits AMPK by disrupting the LKB1 complex, leading to activation of ACC1 and lipogenesis (27). Thus, in addition to its more direct impact on lipogenesis by regulation of lipid metabolic genes (35), our data reveal that AR also supports this metabolic process by activation of 6PGD and the PPP.

In addition to regulation of 6PGD by the androgen signalling axis, our work also revealed that 6PGD can act in a reciprocal manner to maintain AR protein levels and activity. Indeed, S3 was as effective as enzalutamide at inhibiting the expression of some AR target genes, albeit at higher doses. We propose that this positive feedback would serve as an effective circuit to fuel PCa growth and enhance survival. The mechanism(s) by which 6PGD increases AR protein are unknown, although a number of possibilities can be envisioned. First, it has been reported that induction of ROS reduces the levels of AR protein in PCa cells without decreasing its steady state mRNA (36). This post-transcriptional mechanism aligns with our observation that S3/physcion significantly decreased AR protein but only had a negligible impact on AR transcript levels. Second, altered AMPK and SREBP1 signalling downstream of 6PGD/PPP likely influences AR expression. AMPK signalling causes down-regulation of *AR* gene expression as well as promoting AR protein degradation (37). Additionally, SREBP1 has been reported to directly regulate the *AR* gene (38), and one likely consequence of S3-mediated activation of AMPK would be down-regulation of SREBP1. More broadly, the likelihood of shared intermediary factors within each arm of the AR/PPP feedback loop – for example, altered redox homeostasis, SREBP1, AMPK and mTOR – would result in strong positive reinforcement of this complex circuit.

Given the important role of the PPP in PCa growth and survival, established by this study in addition to earlier work (13, 26), targeting this pathway as a possible therapeutic strategy has merit. We investigated this concept using two inhibitors of 6PGD, physcion (1,8-dihydroxy-3-methoxy-6-methyl-anthraquinone; emodin-3-methyl ether) and S3 (1-hydroxy-8-methoxy-Anthraquinone). Physcion (also known as parietin; PubChem CID 10639) was the most active inhibitor of 6PGD activity in an *in vitro* assay amongst a library of ∼2,000 small molecules (27). A plant-derived anthraquinone, physcion was initially investigated for its anti-microbial and anti-inflammatory activities (39). More recently, there has been significant interest in its repurposing as an oncology agent since it has been reported to possess broad anti-cancer activity (i.e. suppression of growth and migration, induction of apoptosis) in leukemia, colorectal, cervical and breast cancer cells, amongst others (27, 40-43). However, while physcion has achieved impressive anti-cancer results in some pre-clinical studies, its poor pharmacological attributes, including low solubility, may impede efforts to progress it to the clinic (39). Therefore, we also tested the physcion derivative compound S3, which has been reported to possess improved pharmacological attributes (27). Our results represent the first evaluation of physcion and S3 in PCa and collectively highlight the potential of therapeutically targeting 6PGD in this disease. Indeed, our data suggest that S3/physcion would possess multi-pronged anti-tumour activity in PCa by: inhibiting oncogenic metabolism, including lipogenesis (i.e. activation of AMPK and suppression of ACC1 and mTOR); increasing levels of ROS, resulting in oxidative stress and lipid peroxidation; and finally, suppressing the levels and activity of AR, the primary oncogenic driver of this disease. Importantly, a Phase I trial reported that physcion was well tolerated with low toxicity (44), supporting its future clinical application.

Since AR-targeted therapies are not curative, there is intense interest in identifying combination therapies that would improve patient outcomes. Our work provides a solid rationale for co-targeting of AR and 6PGD; indeed, we observed synergistic effects of enzalutamide and S3 in PCa models. Moreover, the existence of an AR:6PGD feedback loop enhances the appeal of such a combinatorial strategy. Although we acknowledge that physcion and S3 may not be useful clinical agents due to pharmacological issues, we expect that the future development of therapies that effectively suppress activity of 6PGD, or other components of the PPP, could have a major impact on PCa patients.

## MATERIALS AND METHODS

### Reagents

Chemicals, solvents and solutions, including physcion (C_16_H_12_O_5_; 1,8-dihydroxy-3-methoxy-6-methyl-anthraquinone; emodin-3-methyl ether) and S3 (C_15_H_10_O_4_; 1-hydroxy-8-methoxy-Anthraquinone), were obtained from Sigma-Aldrich (St Louis, MO, USA), except for: enzalutamide (Selleck Chemicals; Houston, TX, USA); apalutamide (ARN-509), darolutamide (ODM-201) and Trolox (Sapphire Bioscience; Redfern, NSW, AUS). All chemicals/reagents were dissolved in dimethyl sulfoxide (DMSO) except dihydrotestosterone (DHT), which was dissolved in ethanol

### Cell line models

LNCaP, VCaP, PC-3 and 22Rv1 human prostate carcinoma cells were obtained from the American Type Culture Collection (ATCC, MD, USA). Dr. Amina Zoubeidi (Vancouver Prostate Centre, Vancouver, Canada) kindly provided LNCaP-V16D (castration-resistant, enzalutamide-sensitive) and LNCaP-MR49F (castration-resistant, enzalutamide-resistant) human prostate cancer cells (28). LNCaP, 22Rv1, V16D and MR49F cells were maintained in RPMI-1640 containing 10% FBS; the media for growth of MR49F cells was additionally supplemented with 10uM enzalutamide. VCaP cells were maintained in Dulbecco’s Modified Eagle’s Medium containing 10% FBS, 1% sodium pyruvate, 1% MEM non–essential amino acids, and 0.1 nM 5α-dihydrotestosterone (DHT). PC-3 cells were maintained in RPMI-1640 containing 5% FBS. All cell lines were authenticated using short tandem repeat profiling in 2018/2019 by ATCC or CellBank Australia.

### Transfection of prostate cancer cell lines

Gene-specific knockdown was achieved by reverse-transfection of PCa cell suspensions (total 5×10^5^ cells) with 12.5 nM siRNA in 6 well plates using RNAiMAX transfection reagent (Life Technologies; Thermo Fisher Scientific, Scornsby, VIC, AUS), according to the manufacturer’s instructions. The siRNAs used in this study were: AR (Silencer Select #4390824/5; s1538, s1539 and custom #4399665; s551824 (Sense: GAACUUCGAAUGAACUACAtt, Antisense: UGUAGUUCAUUCGAAGUUCat,, 6PGD Silencer Select #4427038; s10394 and 10395 and Negative Control 2 #AM4637 (Ambion; Thermo Fisher Scientific) and SREBP1 ON-TARGETplus: 6720 (Dharmacon; Millennium Science, Mulgrave, VIC, AUS).

### Quantitative real-time PCR

Reverse transcription of (1 μg) and qPCR was done as described previously (45). GeNorm (46) was used to identify suitable reference genes: gene expression in cell lines is presented relative to *L19* and *GUSB*, and gene expression in prostate tumour explants is presented relative to *GAPDH, PPIA* and *TUBAIB*. Primers sequences are provided in Table S1.

### Immunoblotting

Whole cell lysates were prepared using RIPA buffer containing cOmplete ULTRA protease and phosphatase inhibitor (Cell Signaling Technology (CST), Danvers, MA, USA) and Western blotting was performed as described previously (47). A list of primary and secondary antibodies used in the study is provided in Table S2.

### RNA sequencing (RNA-seq)

LNCaP cells were seeded at density 5×10^5^ cells in 6-well dishes (Corning) and treated with 1 µM enzalutamide (or 0.1% DMSO control) or transfected with 12.5 nM AR siRNA (or scrambled siRNA control). Each treatment comprised 4 replicates. After 24 h, the cells were collected in Trizol (4 replicates, for RNA analysis) or RIPA Buffer + protease inhibitors (2 replicates, for protein analysis). RNA extractions were completed using RNeasy Mini spin columns (Qiagen, Chadstone, VIC, AUS), according to the manufacturer’s instructions. RNA was eluted in 40 µl RNase-free H_2_O. RT-qPCR and western blotting were performed to verify the expected response of known AR-regulated proteins and genes, PSA/*KLK3* and FKBP51/*FKBP5*. Subsequently, libraries were generated using 800 ng of RNA and NEXTflex Rapid Illumina Directional RNA-Seq Library Prep Kits (Bio Scientific, Kirrawee, NSW, AUS), according to the manufacturer’s instructions. Sequencing was carried out at the South Australian Health and Medical Research Institute Genomics Facility using an Illumina NextSeq 500 (single read 75bp v2 sequencing chemistry). The quality and number of reads for each sample were assessed with FastQC v0.11.3 (48). Adaptors were trimmed from reads, and low-quality bases, with Phred scores < 28, were trimmed from ends of reads, using Trimgalore v0.4.4 (49). Trimmed reads of <20 nucleotides were discarded. Reads passing all quality control steps were aligned to the hg38 assembly of the human genome using TopHat v2.1.1 (50) allowing for up to two mismatches. Reads not uniquely aligned to the genome were discarded. HTSeq-count v0.6.1 (51) was used with the union model to assign uniquely aligned reads to Ensembl Hg38.86-annotated genes. Data were normalized across libraries by the trimmed mean of M-values (TMM) normalization method, implemented in the R v3.5.0, using Bioconductor v3.6 EdgeR v3.20.9 package (52). Only genes expressed at count-per-million value greater than 10 in at least 2 samples per group were retained for further analysis. Differential expressed genes were selected based on the robust version of the quasi-likelihood negative binomial generalized log-linear model (53), with false discovery rate (FDR) set at 0.05. RNA-seq data are available through NCBI’s Gene Expression Omnibus (GSE152254).

### Cell growth and apoptosis assays

Cell growth curves were done using Trypan blue exclusion and manual counting of cells, as described previously (54). Cell viability was also determined by CyQuant™ Assay Cell Proliferation Assays (Thermo Fisher Scientific), according to the manufacturer’s instructions. Apoptosis was measured by collecting cells in FACS binding buffer (47 ml of HANKS buffered saline, 500 µL of Herpes solution and 2.5 mL of 100 mM CaCl_2_), staining with Annexin V PE (BD Pharmagen™, BD Biosciences, CA, US and 1 mM 7-Aminoactiomycin D (Thermo Fisher Scientific) and analysis by Flow Cytometry using a BD LSRFortessa X20.

### Metabolomics

LNCaP cells were seeded at a density of 5×10^5^ cells into Nunclon D multi-dishes with poly lysine coating (Thermo Fisher Scientific), with or without transfection. At time of collection, cells were washed twice with 0.9% w/v NaCl, scraped in MeOH:H2O (1:1).Chloroform was added prior to vortexing, centrifuging and collection of the aqueous layer. The aqueous layer was lyophilised by SpeedVac without heat, then samples were resuspended in 60 µL LC-MS H_2_O and centrifuged at 15,000g at 4°C for 10 min. The supernatants were transferred into HPLC vials, placed at 4°C on the autosampler tray and analysed immediately. Samples were assayed using two different platforms. For the first platform, analyte separation was achieved using a Poroshell 120 HILIC-Z column (2.7 µm, 2.1×100 mm, Agilent) at ambient temperature on a Vanquish-TSQAltis LC-MS/MS system. The pair of buffers used were 95:5 (v/v) water:acetonitrile containing 20 mM ammonium hydroxide and ammonium acetate (Buffer A) and 100% acetonitrile (Buffer B) flowed at 200 µL/min; injection volume was 5 µL. MS acquisition was performed in positive and negative SRM mode to measure amino acids and central carbon metabolites. For the second platform, analyte separation was achieved using a Synergi Hydro-RP column (2.5 µm 3×100mm, Phenomenax) at ambient temperature on a 1260 Infinity (Agilent)-QTRAP500 (AB Sciex) LC-MS/MS system. The pair of buffers used were 95:5 (v/v) water:acetonitrile containing 10 mM tributylamine and 15 mM acetic acid (Buffer A) and 100% acetonitrile (Buffer B) flowed at 200 µL/min; injection volume was 5 µL. MS acquisition was performed in negative SRM mode to measure central carbon metabolites. Calibration standards were injected using the same set up. Raw data was extracted using ProteoWizard and in-house MATLAB scripts.

### Reactive oxygen species (ROS) assays

Cellular ROS levels were measured using CellROX™ Orange Flow Cytometry Assay Kits (Life Technologies). Briefly, 24 h post-seeding (5×10^5^ cells per 6-well plate), the cells were treated with or without antioxidant (0.5 mM Trolox) and left to incubate for the indicated time (siRNA, 48 h; S3,72 h). Cells were stained with CellROX Orange and SYTOX Red Stain and analysed by Flow Cytometry (10-30,000 cells/sample) using a BD LSRFortessa X20.

### Ex vivo *culture of human prostate tumours*

Prostate cancer tissue was obtained with informed written consent through the Australian Prostate Cancer BioResource from men undergoing radical prostatectomy at St Andrew’s Hospital (Adelaide, Australia). Ethical approval for the use of human prostate tumours was obtained from the Ethics Committees of the University of Adelaide (Adelaide, Australia) and St Andrew’s Hospital (Adelaide, Australia). All experiments were performed in accordance with the guidelines of the National Health and Medical Research Council (Australia). The 8 mm core of tissue was dissected and prepared for *ex vivo* culturing as described previously (55). Tissues were treated with AR antagonist 10 µM enzalutamide or 40 µM S3 for 72 h. At the time of collection, the tissues were preserved in RNAlater (Invitrogen; Thermo Fisher Scientific) or formalin-fixed then paraffin embedded.

### Immunohistochemistry (IHC)

Prostate cancer explant tissue sections were evaluated for target antigens 6PGD, Ki67 and pS6 (Ser235/236) by IHC as described previously (55). The antibodies used are shown in Table S2. An automated staining protocol (U OptiView DAB IHC v6 (v1.00.0136)) using the Ventana BenchMark ULTRA IHC/ISH Staining Module (F Hoffmann-La Roche Ltd, Switzerland) was used for the detection of AR. Quantitative image analysis for AR and pS6 (Ser235/236) was completed using FIJI software (ImageJ) (http://fiji.sc/Fiji version 1.52p). Briefly, images (obtained from NDP viewer version 2.7.52; Hamamatsu Photonics K.K, Hamamatsu City, Japan) were imported and converted into three panels using *Colour Deconvolution* plug-in and vector hematoxylin and DAB staining (HDAB) commands. Plug-in *Adjust Threshold* was performed on the DAB-only images to measure % Area (Positivity) and Reciprocal Intensity (R.I). The final DAB intensity values were calculated by subtracting R.I from Maximal Intensity (255) and multiplying by % Area (Positivity). Values from 20-70 images per treatment were measured and R.I was kept constant for each patient.

### Statistical analysis

Data are displayed as the mean; error bars are standard error. Differences between groups were determined using GraphPad Prism with t tests or one-way ANOVA (with Tukey or Dunnett post hoc test), as indicated in the figure legends. A *P* value ≤ 0.05 was considered statistically significant.

## ACKNOWLEDGEMENTS

We acknowledge expert assistance from Kayla Bremert, Madison Helm, Samira Khabbazi, Scott Townley, Zeyad Nassar, Elizabeth Nguyen, Shadrack Mututu, Courtney Moore, Deanna Miller, Mark Van der Hoek, Randall Grose, Bianca Varney and Michelle van Geldermalsen. RNA-seq was performed at the South Australian Health and Medical Research Institute (SAHMRI) Genomics Facility. Flow cytometry analysis was performed at SAHMRI in the ACRF Cellular Imaging and Cytometry Core Facility, which is generously supported by the Australian Cancer Research Foundation, Detmold Hoopman Group and Australian Government through the Zero Childhood Cancer Program. Metabolomics was facilitated by access to Sydney Mass Spectrometry, a core research facility at the University of Sydney. LNCaP-V16D and LNCaP-MR49F cells were a kind gift from Dr. Amina Zoubeidi (Vancouver Prostate Centre, Vancouver, Canada). The authors are grateful to the study participants, as well as the urologists, nurses and histopathologists who assisted in the recruitment and collection of patient information and pathology reports through the Australian Prostate Cancer BioResource.

This work was supported by a grant from Cancer Australia (ID 1138766 to LMB, DJL, MMC, IGM). The research programs of LMB and LAS are supported by the Movember Foundation and the Prostate Cancer Foundation of Australia through a Movember Revolutionary Team Award. LAS and LMB are supported by Principal Cancer Research Fellowships, awarded by Cancer Council’s Beat Cancer project on behalf of its donors, the State Government through the Department of Health and the Australian Government through the Medical Research Future Fund.

---

**Figure S1.**
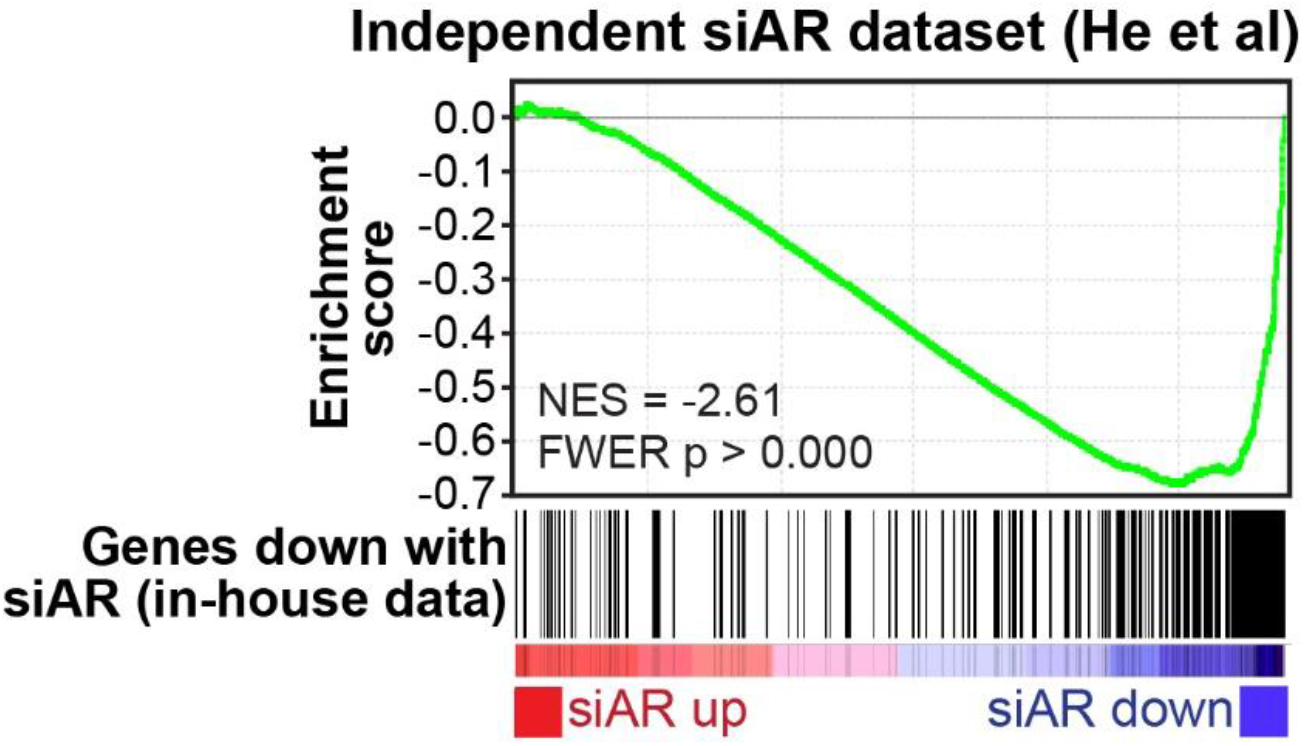
Concordance between our siAR RNA-seq data and an independent dataset, as demonstrated by gene set enrichment analysis (GSEA) (1). RNA-seq data from He and colleagues (2) was kindly provided by Nicholas Mitsiades (Baylor College of Medicine), and genes were ranked by fold-change in siAR treatment versus siControl. Genes down-regulated by siAR versus siControl in our dataset (FDR < 0.01, n = 305) were used as the gene set of interest. Running enrichment scores are plotted (top graph) and normalized enrichment scores (NES) and *P* values are indicated.

**Figure S2.**
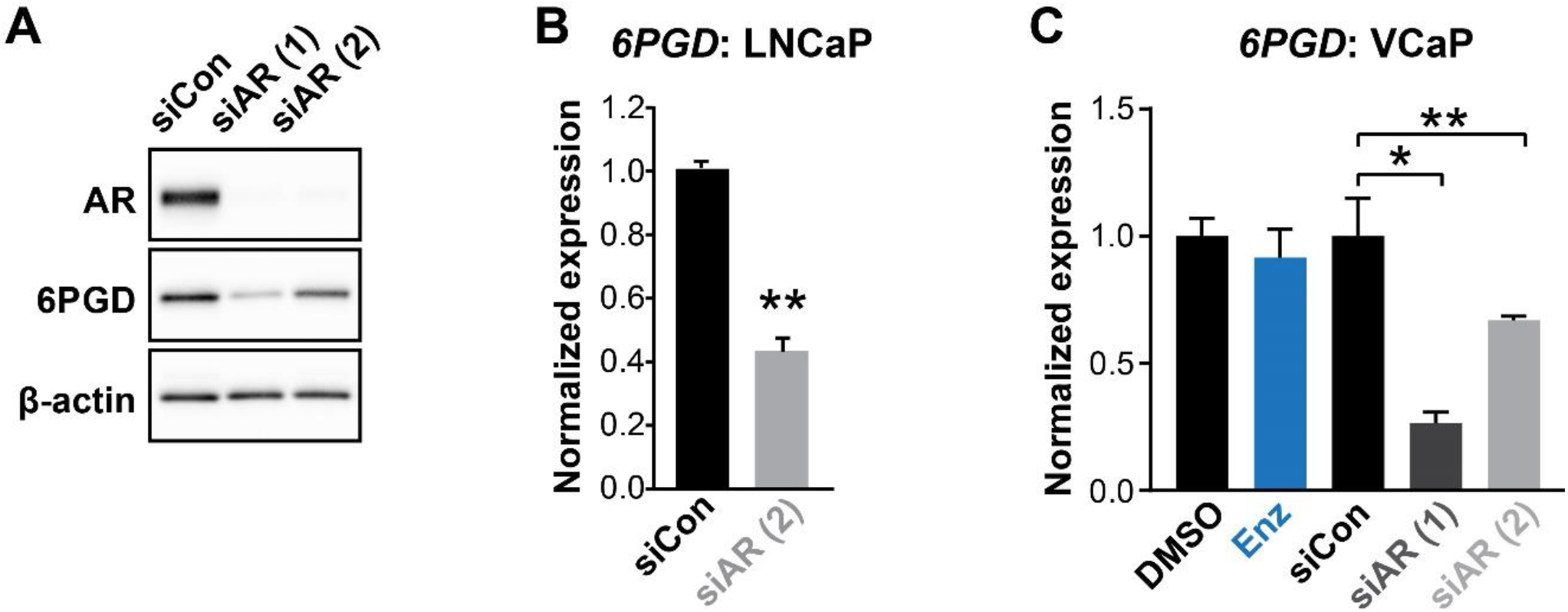
(A) Two distinct AR siRNAs (siAR (1) and siAR (2); 12.5 nM) reduce the expression of 6PGD at the protein level in LNCaP cells. Cells were transfected with 12.5 nM of each siRNA; after 48 h, proteins were extracted and assessed by Western blotting. (B) siAR (2) reduces the expression of 6*PGD* mRNA in LNCaP cells. Transfection of siRNAs was performed as in A. Differential expression was evaluated using an unpaired t test (**, p < 0.001). (C) siAR (1) and siAR (2), but not enzalutamide (Enz, 1 uM), reduce the expression of 6*PGD* mRNA in VCaP cells. Transfection of siRNAs was performed as in A. Cells were treated with DMSO or Enz for 24 h. Differential expression compared to DMSO or siCon was determined using ANOVA and Dunnett’s multiple comparison tests (*, p < 0.05; **, p < 0.01).

**Figure S3.**
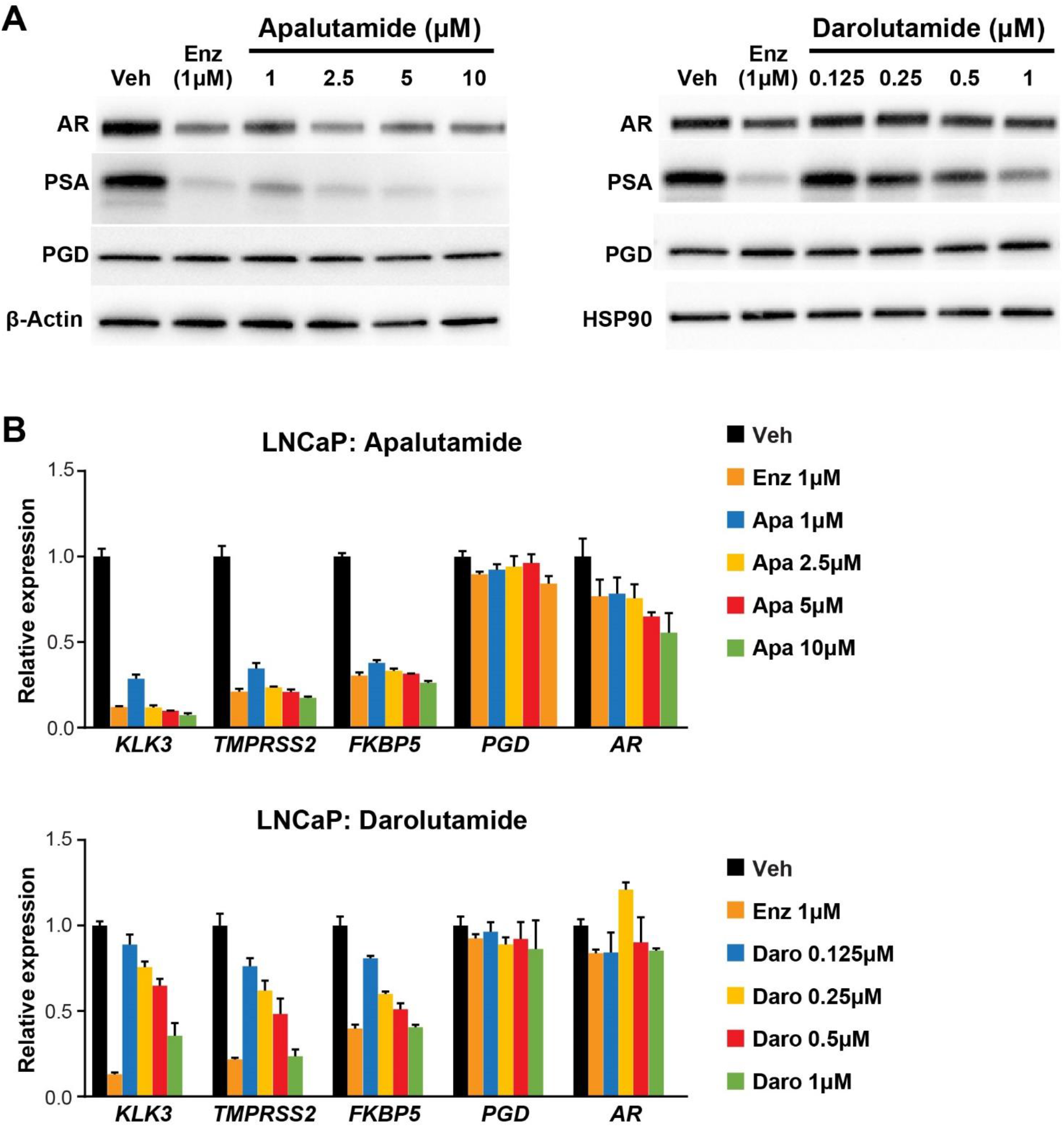
Next-generation AR antagonists apalutamide and darolutamide inhibit AR target gene expression at the protein (A) and mRNA (B) level, but do not reduce expression of 6PGD protein or mRNA. Cells were treated for 24 h with the indicated doses of each drug.

**Figure S4.**
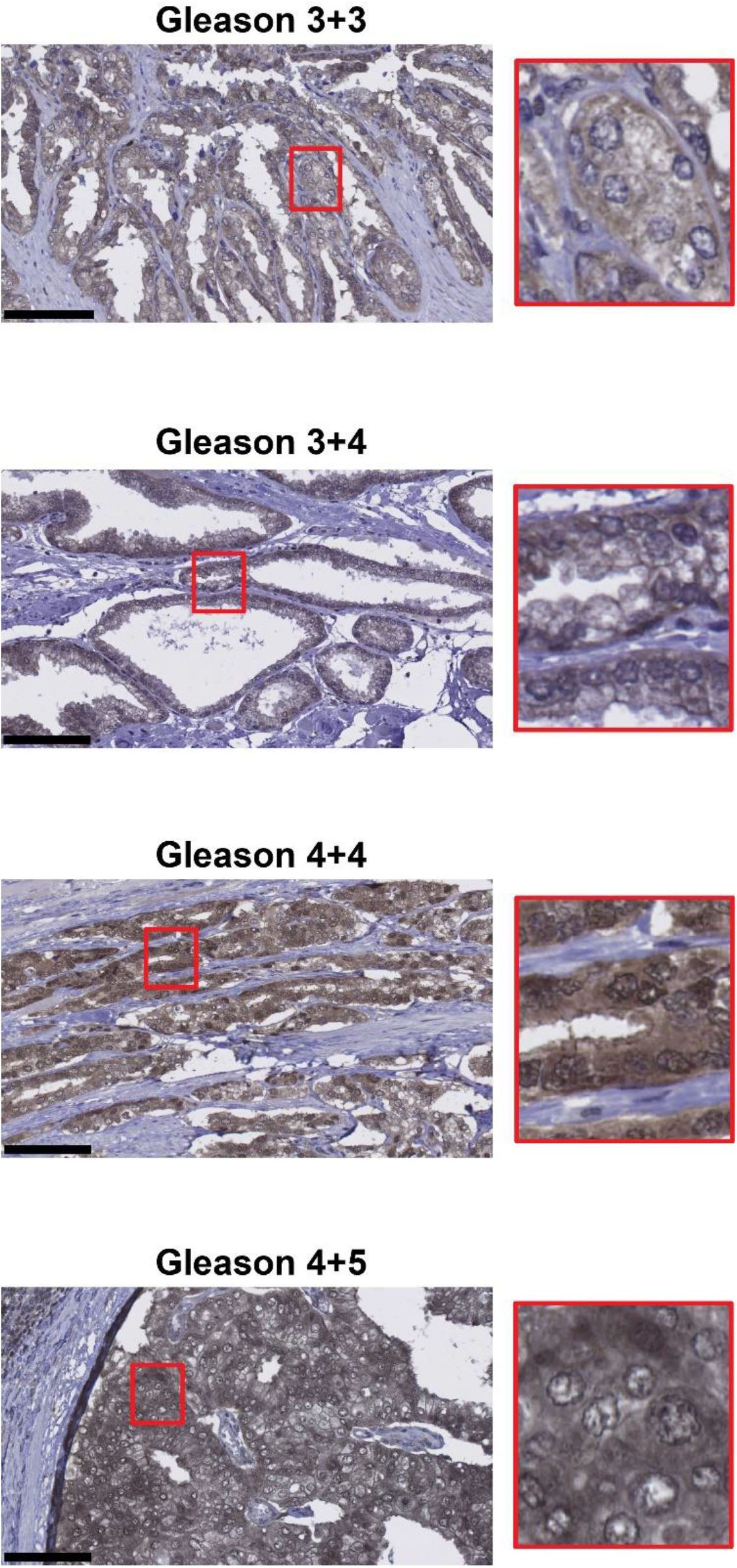
Representative images of 6PGD IHC in patient tumours. Gleason grades are shown. Scale bars represent 100 µm.

**Figure S5.**
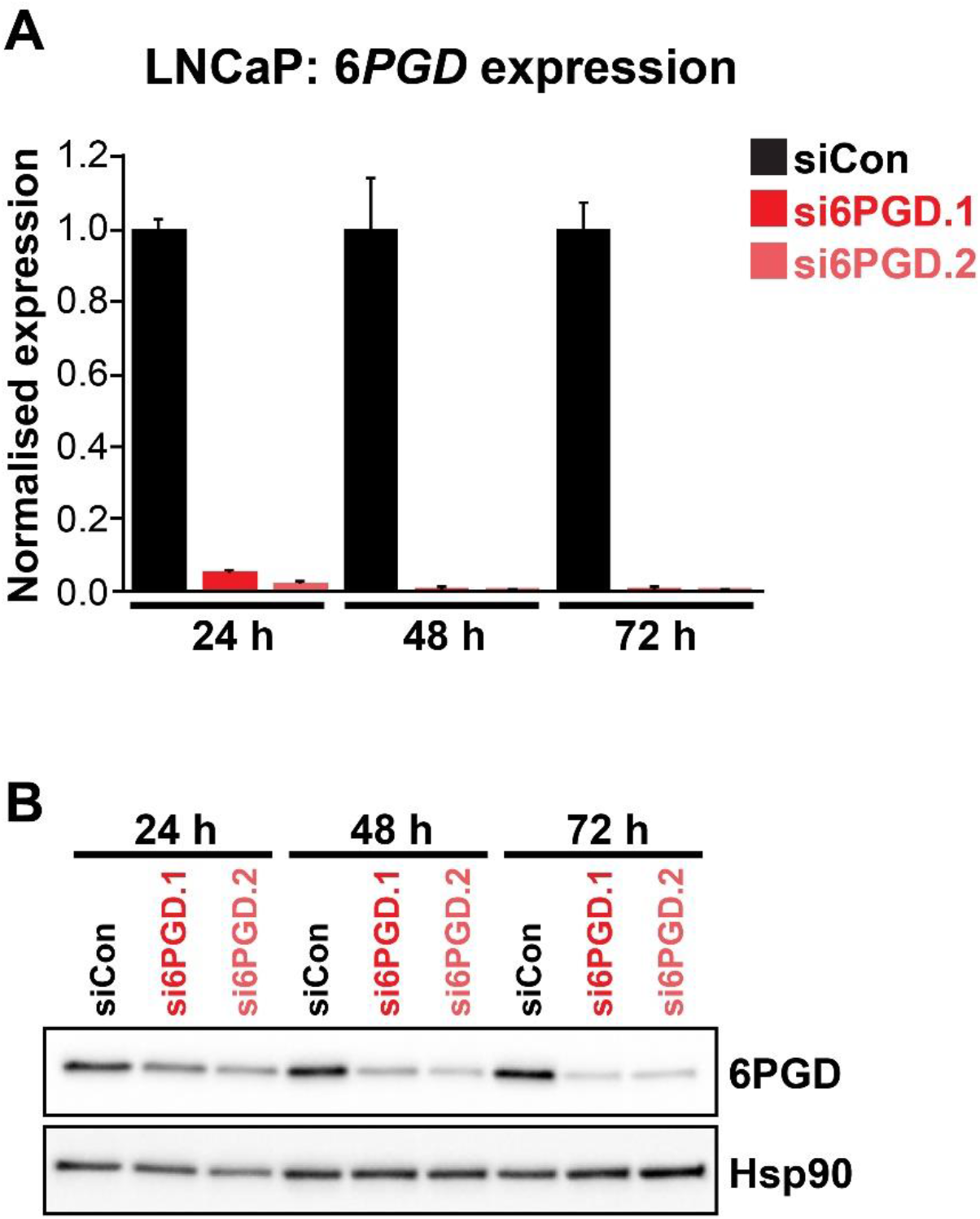
Two distinct 6PGD siRNAs (si6PGD.1 and si6PGD.2) effectively reduce 6PGD expression in LNCaP cells. Cells were transfected with 12.5 nM of each siRNA for 72 h, after which 6PGD mRNA was measured by RT-qPCR (A) or 6PGD protein was measured by immunoblotting (B).

**Figure S6.**
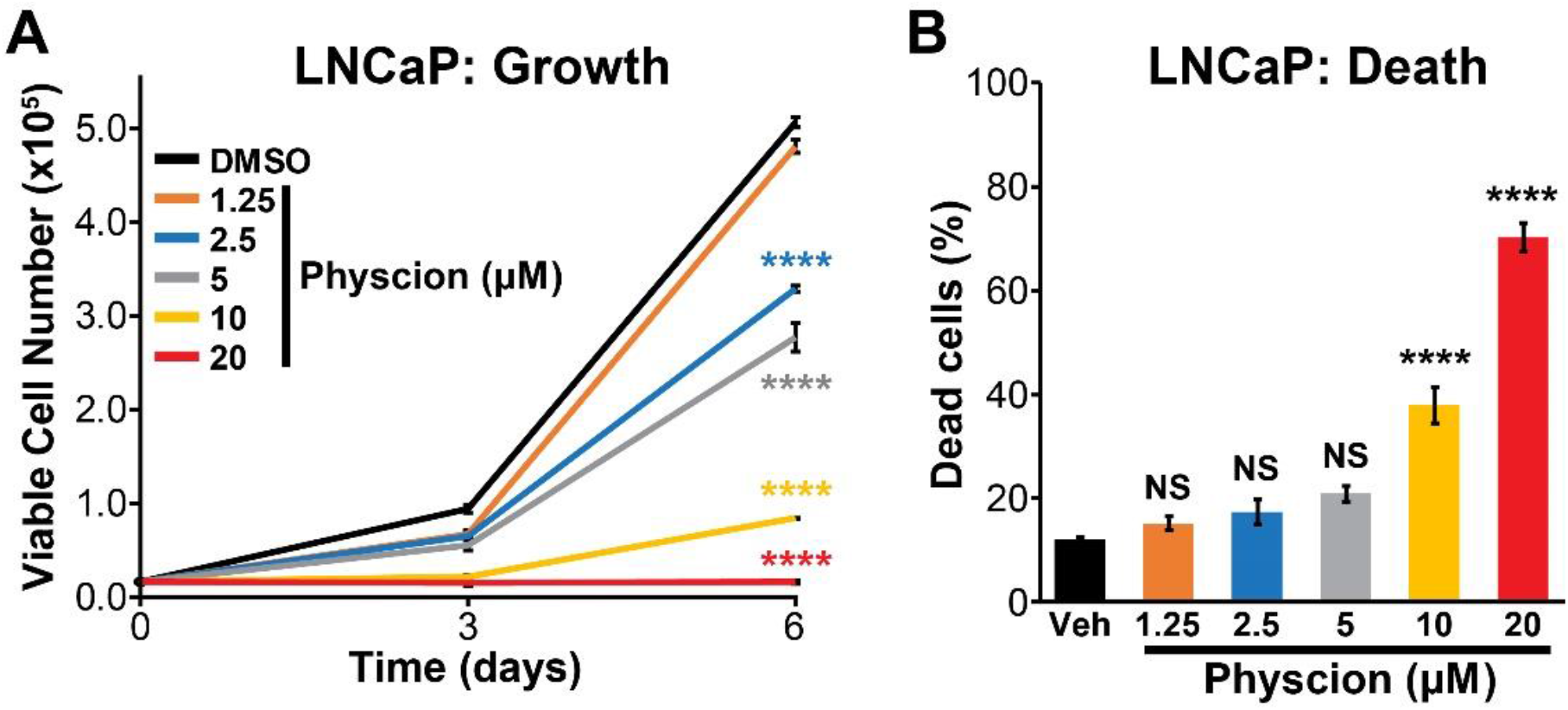
Physcion effectively suppresses growth (A) and causes death (B) of LNCaP cells. Live and dead cells were measured (A, at the indicated time-points; B, at day 6) using Trypan blue exclusion assays. Physion’s effects on growth and death compared to vehicle (Veh) were determined using ANOVA and Dunnett’s multiple comparison tests (****, p < 0.0001; NS, not significant).

**Figure S7.**
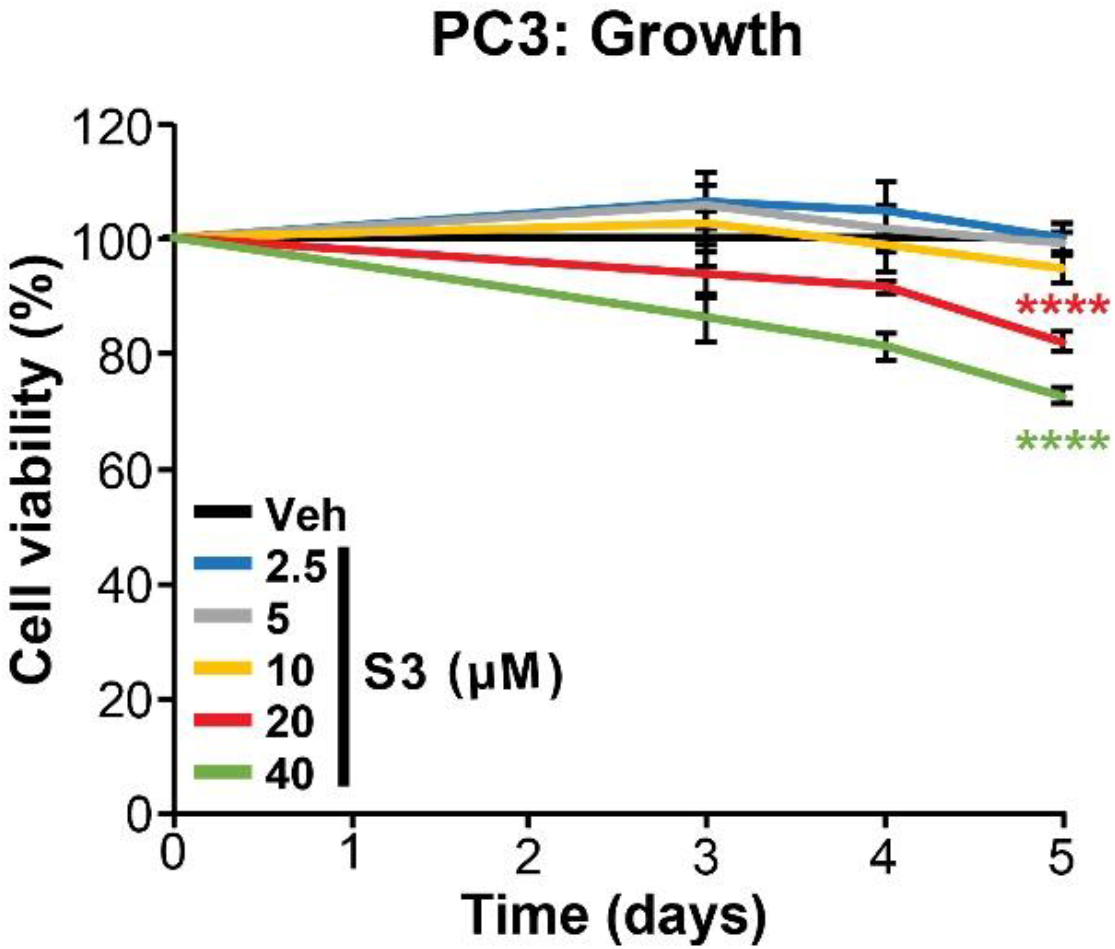
Effect of S3 on growth of PC3 cells. Cell viability assessed by CyQuant Direct Cell Proliferation Assay. Fluorescence at day 0 was set to 100%. The effect of S3 on growth compared to vehicle (Veh) was determined using ANOVA and Dunnett’s multiple comparison tests; only 20µM and 40µM doses were significantly different to Veh (****, p < 0.0001).

**Figure S8.**
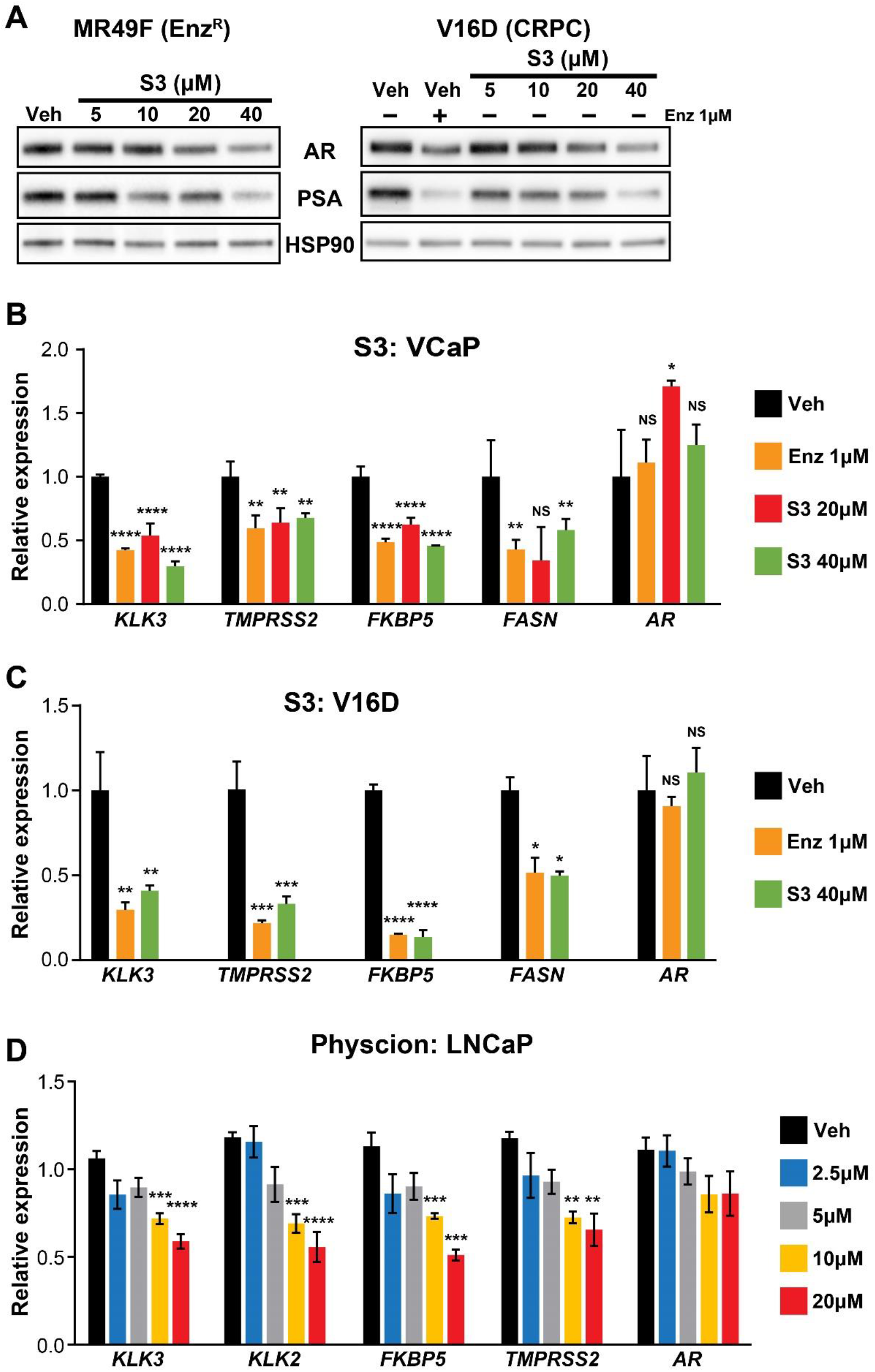
(A) S3 decreases AR and PSA protein levels in MR49F (left) and V16D (right) cells. Protein was extracted from cells at 24h and assessed by Western blotting. HSP90 is shown as a loading control. (B-C) S3 suppresses AR target gene expression in VCaP (B) and V16D (C) cells after 24 h treatment. Expression is shown relative to *GUSB* and *L19*; vehicle (Veh) was set to 1. (D) Physcion suppresses AR target gene expression in LNCaP cells after 24 h treatment. Expression is shown relative to *GUSB* and *L19*. Differential expression compared to vehicle (B-D) was determined using ANOVA and Dunnett’s multiple comparison tests (*, p < 0.05; **, p < 0.01; ***, p < 0.001; ****, p < 0.0001; NS, not significant).

